# The effects of long duration spaceflight on sensorimotor control and cognition

**DOI:** 10.1101/2021.06.22.449414

**Authors:** GD Tays, KE Hupfeld, HR McGregor, AP Salazar, YE De Dios, NE Beltran, PA Reuter-Lorenz, IG Kofman, SJ Wood, JJ Bloomberg, AP Mulavara, RD Seidler

## Abstract

Astronauts returning from spaceflight typically show transient declines in mobility and balance. These whole-body postural control behaviors have been investigated thoroughly, while study of the effects of spaceflight on other sensorimotor behaviors is prevalent. Here, we tested the effects of the spaceflight environment of microgravity on various sensorimotor and cognitive tasks during and after missions to the International Space Station (ISS). We obtained mobility (Functional Mobility Test), balance (Sensory Organization Test-5), bimanual coordination (bimanual Purdue Pegboard), cognitive-motor dual-tasking and various cognitive measures (Digit Symbol Substitution Test, Cube Rotation, Card Rotation, Rod and Frame Test) before, during and after 15 astronauts completed 6+ month missions aboard the ISS. We used linear mixed effect models to analyze performance changes due to entering the microgravity environment, behavioral adaptations aboard the ISS and subsequent recovery from microgravity. We identified declines in mobility and balance from pre- to post-flight, suggesting possible disruption and/or downweighting of vestibular inputs; these behaviors recovered to baseline levels within 30 days post-flight. We also identified bimanual coordination declines from pre- to post-flight and recovery to baseline levels within 30 days post-flight. There were no changes in dual-task performance during or following spaceflight. Cube rotation response time significantly improved from pre- to post-flight, suggestive of practice effects. There was a trend for better in-flight cube rotation performance on the ISS when crewmembers had their feet in foot loops on the “floor” throughout the task. This suggests that tactile inputs to the foot sole aided orientation. Overall, these results suggest that sensory reweighting due to the microgravity environment of spaceflight affected sensorimotor performance, while cognitive performance was maintained. A shift from exocentric (gravity) spatial references on Earth towards an egocentric spatial reference may also occur aboard the ISS. Upon return to Earth, microgravity adaptions become maladaptive for certain postural tasks, resulting in transient sensorimotor performance declines that recover within 30 days.

## 1. Introduction

There are well-documented changes in human sensorimotor performance following spaceflight, including post-flight declines in locomotion, balance, and fine motor control (Thornton & Rummel, 1977; Paloski et al., 1992; 1994; Reschke et al., 1994a; 1994b; 1998; Black et al., 1995; Mcdonald et al. 1996, Bloomberg et al. 1997; Newman et al. 1997; Layne et al. 1998; 1999; Bock et al., 2003; Campbell et al., 2005; Rafiq et al. 2006). However, the effects of spaceflight on human cognition and other motor behaviors have not been as thoroughly investigated (Strangman et al., 2014; Garrett-Bakelman et al., 2019). Performance of whole-body postural control typically returns to pre-flight levels within approximately two weeks of return to Earth (Wood et al., 2015; Ozdemir et al., 2018), however it is not clear whether the same is true for other sensorimotor or cognitive behaviors.

Vestibular inputs are altered during spaceflight; in particular, otolith (the small structure within the inner ear that senses linear accelerations and tilt) signaling of head tilt, which relies upon gravity is absent and likely gets down weighted (Reschke et al., 1994a; 1994b; 1998; Paloski et al., 1992; 1994; Black et al., 1995; 1999; Clément et al., 2020). The central nervous system adapts to altered vestibular inputs in-flight due to microgravity with as little as 2 weeks spent in spaceflight (Layne et al., 1998). Upon return to Earth however, these adaptive changes may become maladaptive, resulting in difficulties with whole-body motor control. Post-flight impairments in locomotion (Mulavara et al., 2018; Miller et al., 2018; Layne et al., 1998; Mcdonald et al., 1996; Bloomberg et al., 1997), balance (Reschke et al., 1994a; 1994b; 1998; Paloski et al., 1992; 1994; Black et al., 1995; 1999), jumping (Newman et al. 1997), obstacle navigation (Mulavara et al., 2010; Bloomberg et al. 2015), and eye-head coordination (Reschke et al., 2017) have been reported. Neural processing and motor control re-adapt to the presence of Earth’s gravity in the weeks following return, with performance returning to pre-flight levels with about 6 days on a variety of functional tasks (Miller et al., 2018) to 15 days for the functional mobility test (FMT; Mulavara et al., 2010).

In-flight changes in performance of fine motor tasks have also been identified. For instance, astronauts maintained their manual dexterity while performing survival surgery on rats during a Neurolab shuttle mission. However, there was a significant increase in operative time, in some cases taking 1.5 to 2 times longer than on Earth (Campbell et al., 2005), which may be indicative of a speed-accuracy trade-off. Indices of movement variability, reaction time, and movement duration also increased on a hand pointing task executed without visual feedback during Neurolab Space Shuttle missions (Bock et al., 2003), in addition to a significant increase in movement amplitude shortly following landing. During Skylab missions, impairments in reaching and grasping were also documented (Thornton & Rummel, 1977). Additionally, decreases in both force regulation and performance quality while tying surgical knots were identified in the low gravity phase of parabolic flight (Rafiq et al. 2006). Recently, it has been shown that long duration spaceflight results in decreases in fine motor control, as seen by an increase in completion time on a grooved pegboard test (Mulavara et al., 2018). Here we evaluate bimanual motor coordination pre- and post-flight using the bimanual Purdue Pegboard Test.

Several spaceflight stressors have the potential to impact cognition in-flight, including sleep loss, motion sickness, and social isolation. Astronauts anecdotally report so-called “space fog”, including attention lapses, short term memory problems, confusion, and psychomotor problems (Clément et al. 2020). Previous investigations into characterizing the cognitive effects of spaceflight have failed to strongly support or refute such effects (c.f. Strangman et al., 2014). One study showed, however, an increased ability to mentally rotate the visual image of their environment as their exposure to microgravity increased, yet also a decreased ability in spatial orientation of written letters during the first 5 days in-flight (Clément et al., 1987). They posit that the disappearance of a reference field (e.g. the ground) may affect the central representation of movements.

There have also been reported declines in the ability to perform simultaneous cognitive and motor dual-tasking in-flight (Manzey et al. 1995; 1998). The authors suggested that an increased demand for cognitive control of movement in microgravity may interfere with simultaneous cognitive task performance. This was further supported by Bock et al. (2010), who found higher tracking error inflight in both the single and dual-task conditions and higher dual-task cost in a rhythm production reaction-time task compared to a visuospatial reaction-time task and a choice reaction-time task. The authors suggest that this may be due to a scarcity of resources required for complex motor programming due to sensorimotor adaptation to microgravity. Dual-tasking deficits in astronauts post-flight were also identified when astronauts were measured in a tracking task whilst responding and entering numerical codes with their non-dominant hand (Moore et al., 2019). In addition, NASA’s “twins study” also showed increased risk-taking on a cognitive task throughout spaceflight, as well as decreased accuracy in a visual object learning task, decreased abstract shape matching, and decreased cognitive speed for all measures on a subset of tasks from the Penn Computer Neurocognitive Battery, except for the digit symbol substitution task post-flight (Garrett-Bakelman et al., 2019). However, the twins study only tested one astronaut in-flight and compared performance to that of their Earth bound twin, and other previous investigations similarly had small sample sizes (Manzey et al., 1995; 1998; Bock et al., 2010). It remains unclear whether or how cognitive function is impacted by spaceflight. Spaceflight analog environments, such as extended isolation, reduced spatial cognition (Stahn et al., 2019) and head-down tilt bedrest (HDBR) have shown to result in an overall cognitive slowing (Basner et al., 2021). Moreover, spatial orientation and distance estimation are impaired by both the hypergravity and microgravity phases of parabolic flight (Clément et al., 2016). Thus, here we also evaluated performance on a range of cognitive assessments pre- and post-flight.

As NASA’s goal shifts from the International Space Station (ISS) to the Moon and Mars, mission duration will increase. It is imperative to understand how other factors may interact with microgravity to affect sensorimotor and cognitive function, particularly flight duration, age and sex. Exploration missions to Mars surface are estimated to take around 30 months in total (Clément et al., 2019), making it important to understand how mission duration interacts with changes in sensorimotor and cognitive function with spaceflight. Associations between mission duration and the magnitude of brain structural changes, free water shifts, and ventricular enlargement have been previously reported (Hupfeld et al., 2020a; Roberts et al., 2017; Alperin 2017). There is also evidence that longer flight duration results in prolonged brain and behavior recovery profiles (Bryanov et al. 1976; Hupfeld et al. 2020a). Flight duration may also be correlated with the magnitude of sensorimotor and cognitive changes that occur with spaceflight, or that effects of flight duration may be due to an interaction of microgravity with isolation and confinement hazards.

As age increases, sensorimotor adaptability declines (Seidler et al., 2010; Anguera et al., 2011). Astronaut training requires years to complete and the average age for an astronaut at the onset of their first mission is 39.8 (±5.28) years (Smith et al., 2020). It is important to consider the impact of age on behavioral and brain changes in spaceflight, thus we include age as a model covariate for exploratory purposes. Sex differences in the effects of microgravity have rarely been considered (as the Astronaut Corps has been historically male (Reschke et al., 2014)), but with the future Artemis program having equal representation of the sexes, it is imperative to identify any sex related differences. While our sample size of 15 astronauts is not large enough for a well-powered investigation of sex effects, we include sex as a model covariate for exploratory purposes.

Here we aimed to investigate the spaceflight impacts on sensorimotor and cognitive performance. We included several assessments of whole-body sensorimotor behaviors including the Functional Mobility Test (FMT) and Sensory Organization Test-5 (SOT-5) implemented using computerized dynamic posturography. We also assessed fine motor control using the bimanual Purdue Pegboard Test. Finally, we assayed multiple aspects of cognitive function including processing speed, mental rotation, spatial working memory and cognitive-motor dual-tasking. Most tests were administered pre- and post-flight, with a subset of the test battery performed on three occasions on the ISS. Follow-up performance measurements were obtained over six months post-flight to characterize the trajectory of re-adaptation following return to Earth.

We hypothesized, based on prior trajectories of change (Mulavara et al., 2010; Wood et al., 2015), performance on all sensorimotor tasks would decline from pre- to post-flight, and then recover to pre-flight levels within one month. We further hypothesized that performance on cognitive tasks would decrease from pre- to post-flight, with a similar recovery profile as sensorimotor tasks. Finally, we hypothesized that astronauts’ sensorimotor and cognitive (dual-tasking, spatial working memory) performance would be disrupted following their arrival to the ISS, and would then resolve throughout the flight as they adapted to microgravity.

## 2. Materials & Methods

### 2.1 Participants

Fifteen astronauts participated in this study (Table 1). One withdrew from the study prior to their last post-flight data point. The mean age at launch in this study was 47.46 years (± 6.28). 26% of the participants were female. Mission duration to the ISS lasted an average of 188.13 days (± 57.46). Six astronauts had previous flight experience, having spent an average of 75 days (± 131.36) in space across an average of 0.8 (± 1.15) previous missions. An average of 5.77 years (± 1.6) had elapsed since the end of their previous mission. The University of Michigan, University of Florida, and NASA Institutional Review Boards approved all study procedures. All participants provided their written informed consent. This study was implemented as part of a larger NASA-funded project (NASA #NNX11AR02G) aiming to investigate the extent, longevity, and neural bases of long-duration spaceflight-induced changes in sensorimotor and cognitive performance (Koppelmans et al., 2013).

**Table 1:**
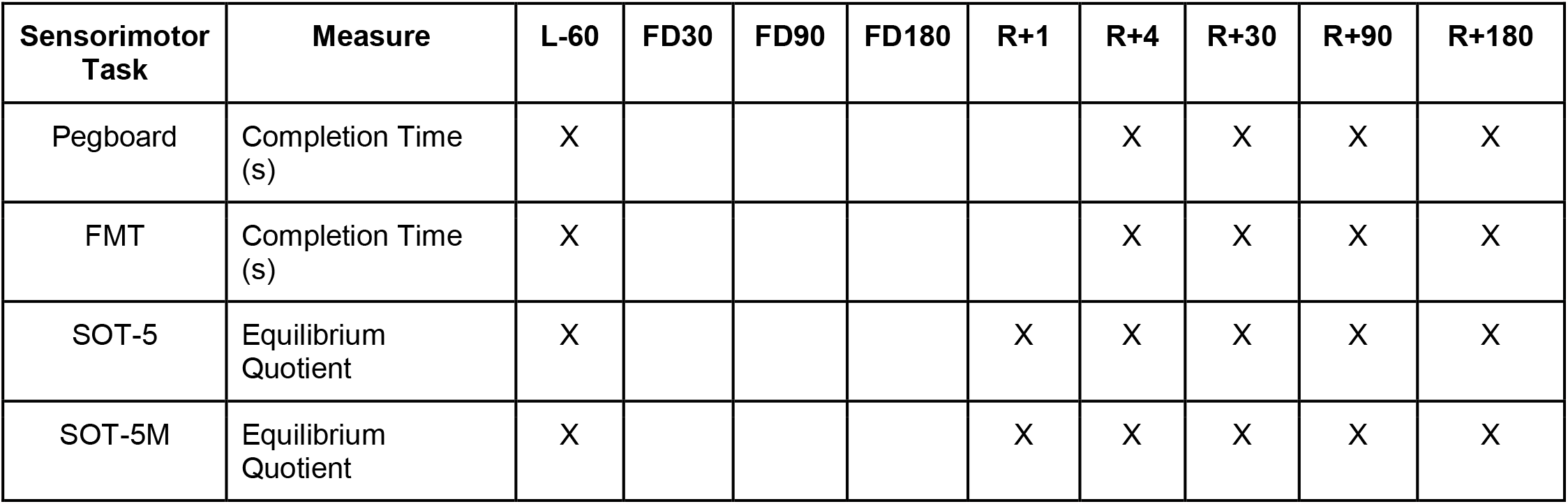
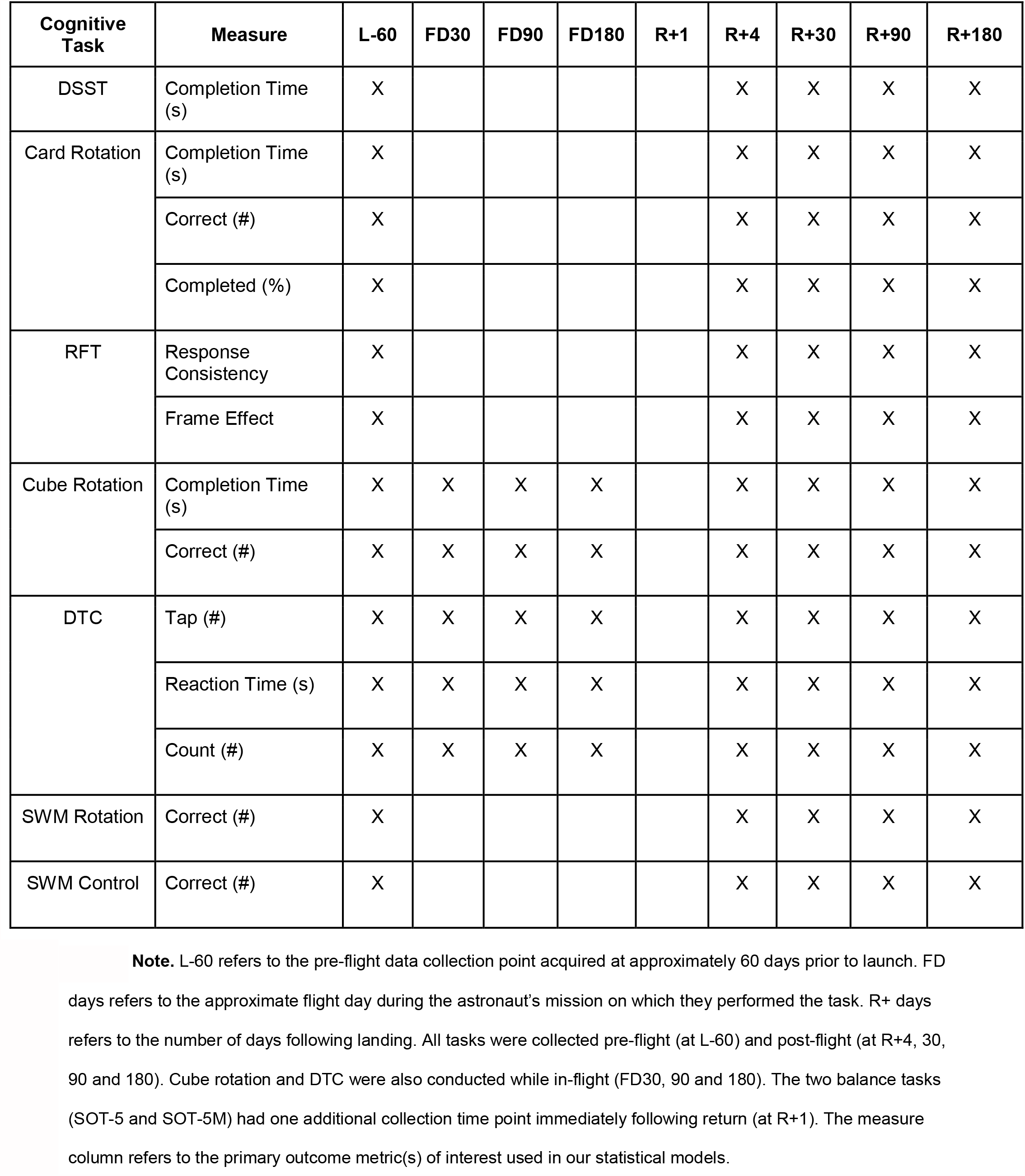
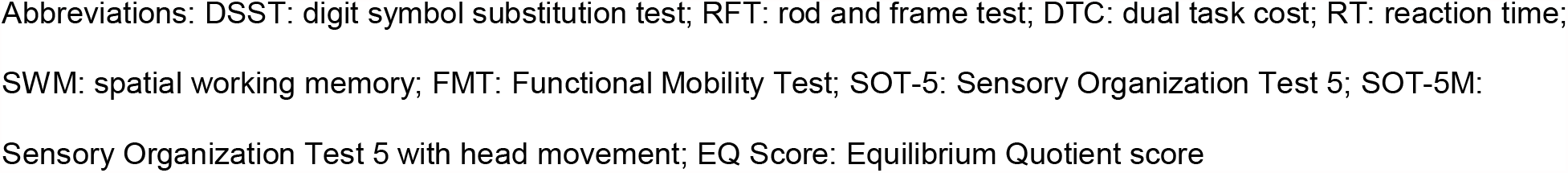
Tasks and data collection time points

### 2.2 Behavioral Assessments

#### 2.2.1 Sensorimotor Measures

##### 2.2.1.1. Whole-body postural and locomotor control

To assess performance changes in relation to spaceflight for whole-body postural control, we administered several balance and locomotion tests. We used the Functional Mobility Test (FMT; Mulavara et al. 2010) to assess ambulatory mobility. This test was designed to assess movements similar to those required during spacecraft egress, which are measured by total completion time. The FMT is a 6 × 4 m obstacle course that requires participants to step over, under and around foam obstacles and change heading direction. Participants start from an upright seated position, buckled into a 5-point harness. After releasing their harness and standing up, they walked on a firm surface for the first half of the test and on a medium density foam for the second half. This compliant foam makes surface support and proprioceptive inputs unreliable (Mulavara et al., 2010). Astronauts performed the FMT 10 times as quickly as possible. For analysis purposes we only analyzed completion time on the first trial to minimize the effects of task learning.

Dynamic postural control was assessed using Computerized Dynamic Posturography (Equitest, NeuroCom International, Clackamas, OR; Reschke et al., 2009). Specifically, astronauts completed the Sensory Organization Test-5 (SOT-5 and SOT-5M). We administered SOT-5, in which the eyes are closed and the platform is sway-referenced, forcing greater reliance on vestibular inputs. We also administered SOT-5M, in which participants make ±20° head pitch movements at 0.33 Hz paced by an auditory tone (Wood et al., 2012). At each pre- and post-flight time point, we administered three trials of the SOT-5 and SOT-5M. Equilibrium Quotient scores were derived from peak-to-peak excursion of the center of mass (estimated at 55% of total height) over a 20-second trial (Nashner, 1972; Paloski et al., 1992). As in our previous work, we used the median Equilibrium Quotient score from each time point in all statistical analyses (Lee et al., 2019).

##### 2.2.1.2. Fine motor control

To assess bimanual coordination, we used the bimanual Purdue Pegboard Test (Tiffin & Asher, 1948). The bimanual Purdue Pegboard Test is a well validated measure of bimanual manual dexterity. Participants were instructed to place 15 small metal pegs into fitted holes. We used their completion time to place all the pegs with both hands for statistical analysis.

#### 2.2.2. Cognitive measures

##### 2.2.2.1. Cognitive-Motor Dual-Tasking

We assessed dual-tasking using a motor and a cognitive task, both separately and simultaneously. The motor task required the participant to perform a choice button press, of two possible buttons, when an “X” was displayed in one of two boxes positioned on either side of the computer screen, cueing the participants to press the button on the corresponding side. The cognitive task required participants to monitor a separate box, positioned directly above the button press indicating boxes, that rapidly changed colors and to count the number of times that the box turned blue (this occurred infrequently relative to other colors, making this akin to an oddball detection task). Each task was performed alone in a single task (ST) conditions as well as together in a dual-task (DT) condition. Performance declines between single to dual-task conditions are frequently referred to as dual-task cost (DTC). DTC has been shown to be a marker of resource limitation for task performance (Tombu & Joliocoeur, 2003) and served as our performance metric, calculated as the change when dual-tasking relative to single tasking ((*DT* - *ST*)/*ST* ∗ 100). Higher DTC during spaceflight would suggest more interference and higher processing loads. We have previously used this task to analyze dual-tasking changes in HDBR analog environments (Yuan et al., 2016).

##### 2.2.2.2. Spatial working memory

We used three task to asses spatial working memory; 1) a spatial working memory task (SWM; Anguera et al., 2010), 2) Thurstone’s 2D card rotation test (Ekstrom et al., 1976) and 3) three-dimensional cube figure mental rotation task (Shepard & Metzler, 1988). During the SWM task, participants were instructed to mentally connect three dots that formed the points of a triangle. Then, after a 3000ms retention phase three new dots would appear on the screen and the participant must decide if they form the same triangle, but rotated, or a different triangle. Participants also performed a control task in which they were shown three dots forming a triangle and then, following a 500ms retention phase, one dot appeared and they would identify if that dot was one of the original three (Anguera et al., 2010; Salazar et al., 2020). We collected 30 trials of each task. For both tests, we used the response time and number of correct responses as our outcome measures. During the 2D card rotation task, participants first were presented with a 2D drawing of an abstract shape. Then they were presented with another drawing and were instructed to identify if it was the same shape rotated or a different shape (the original shape mirrored or altogether different) (Ekstrom et al., 1976; Salazar et al., 2020). The completion time, amount completed and accuracy were recorded and utilized for analyses. Finally, the cube rotation task had participants observe a 3D cube assembly for 3 seconds. Following a 2 second retention phase, two new cube assemblies would appear on the screen and the participant was instructed to identify which of the two matched the initial target image (Shepard & Metzler, 1988; Salazar et al., 2020). Reaction time and accuracy were analyzed for this task. The 3D cube rotation task was administered twice per session while in spaceflight; it was first performed with participants free floating in microgravity (referred to as Cube 1, tethered to a workstation), then with the crewmember in a posture that mimics a seated position with the feet on the “floor” in foot loops (referred to as Cube 2).

##### 2.2.2.3. Rod and Frame Test

Visual field dependence was assessed with the Rod and Frame Test (RFT); in which the participant looks into a “tunnel” (to remove peripheral visual cues) and attempts to align a rod to Earth vertical. This has been shown to identify visual-vestibular interactions (Witkin & Asch, 1948). Outcome measures for the RFT were frame effect, measured as the angular deviation between the participants perceived vertical and true vertical, and the response consistency (sometimes referred to as response “variability”, although in the present work we will refer to this metric as response consistency).

##### 2.2.2.4. Digit Symbol Substitution Task

We utilized the Digit Symbol Substitution Test (DSST) to analyze cognitive processing speed. During this task the participants were presented with a sheet of paper that required them to match numbers with symbols according to a key that is provided at the top of the page (Weschler, 1986). We measured completion time and the number correct as outcome measures for analysis.

### 2.3 Testing Timeline

As shown in Figure 1, astronauts performed all behavioral tasks prior to launch (180 and 60 days pre-flight), and four times following their return to Earth (approximately 4, 30, 90, and 180 days post-flight);. The initial testing point of 180 days before launch (L-180) was used as a familiarization session and was not included in the analyses here. Sensorimotor (FMT, SOT-5, SOT-5M and Bimanual Pegboard) and cognitive (DSST, Card Rotation, RFT, SWM, and dual-tasking) tasks were all measured 60 days before flight and then within a few days of returning to Earth to elucidate the effects of long-term microgravity exposure. SOT-5 and SOT-5M data had an additional data collection time point approximately one day following post-flight. These same measures were all recorded over the following six months post-flight to allow us to investigate recovery from any performance changes due to spaceflight and the microgravity environment. Post-flight testing sessions occurred between 1 to 7 days after landing; to account for this, the time difference between landing and the first post-flight time point was used as a model covariate in analyses. In addition, a subset of tasks (cube rotation and dual-tasking) were collected three times during spaceflight (FD (Flight Day) 30, FD90 and FD150); this allowed us to determine the direct effects of microgravity on performance of these tasks.

**Figure 1.**
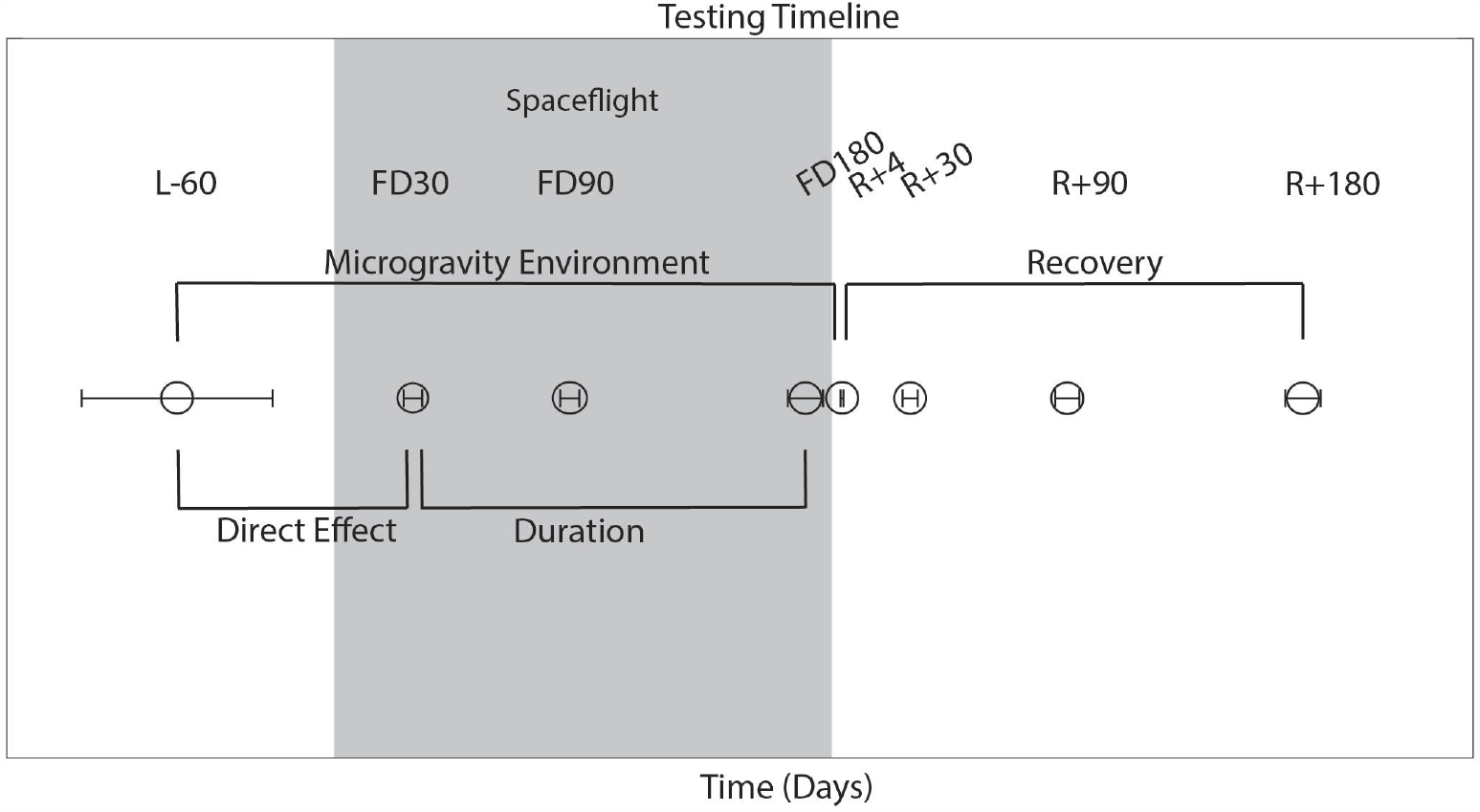
Testing Timeline. L: Launch, R: Return, FD: Flight Day, time spent during spaceflight. Launch occurred on day 0. The average day of data collection is plotted relative to launch, with error bars indicating standard deviation.

#### 2.2.3. Statistical Analyses

We used the nlme package (Pinheiro et al., 2020) in R 3.6.1 (R Core Team, 2019) to fit linear mixed effects models with restricted maximum likelihood (REML) estimation for performance changes over time. In each model subject we entered subject as a random intercept to allow for different starting points for each person (as in our previous work Koppelmans et al., 2017). Specifically, our first model evaluated the effect of the microgravity environment, testing for pre-flight (L-60) to post-flight (R+1/R+4) changes. Our second model evaluated the recovery from the microgravity environment, testing for changes across post-flight time points (R+1/R+4, R+30, R+90, R+180) in measures that showed significant change pre- to post-flight. Our third model evaluated the direct effects of microgravity, testing for performance changes from pre-flight (L-60) to the first in-flight test day (FD30). Our final model evaluated the effects of duration aboard the ISS, testing for changes in performance across the three in-flight test sessions (FD30, FD90, FD150) on select measures. For 3 of the 17 (Card rotation completed, Tap DTC & Count DTC) measures analyzed, the residuals were not normally distributed. We addressed this by log transforming the data prior to statistical analyses (Ives, 2015), however for these three measures transformation did not normally distribute the residuals. Given this, the results of the Card rotation number completed, Tap DTC & Count DTC measures should be interpreted with caution. To account for multiple comparisons, we corrected *p*-values within each of the models using the Benjamini-Hochberg false discovery rate (FDR) correction (Benjamini & Hochberg, 1995); we present the FDR-corrected *p*-values in Tables 2-5.

**Table 2:**
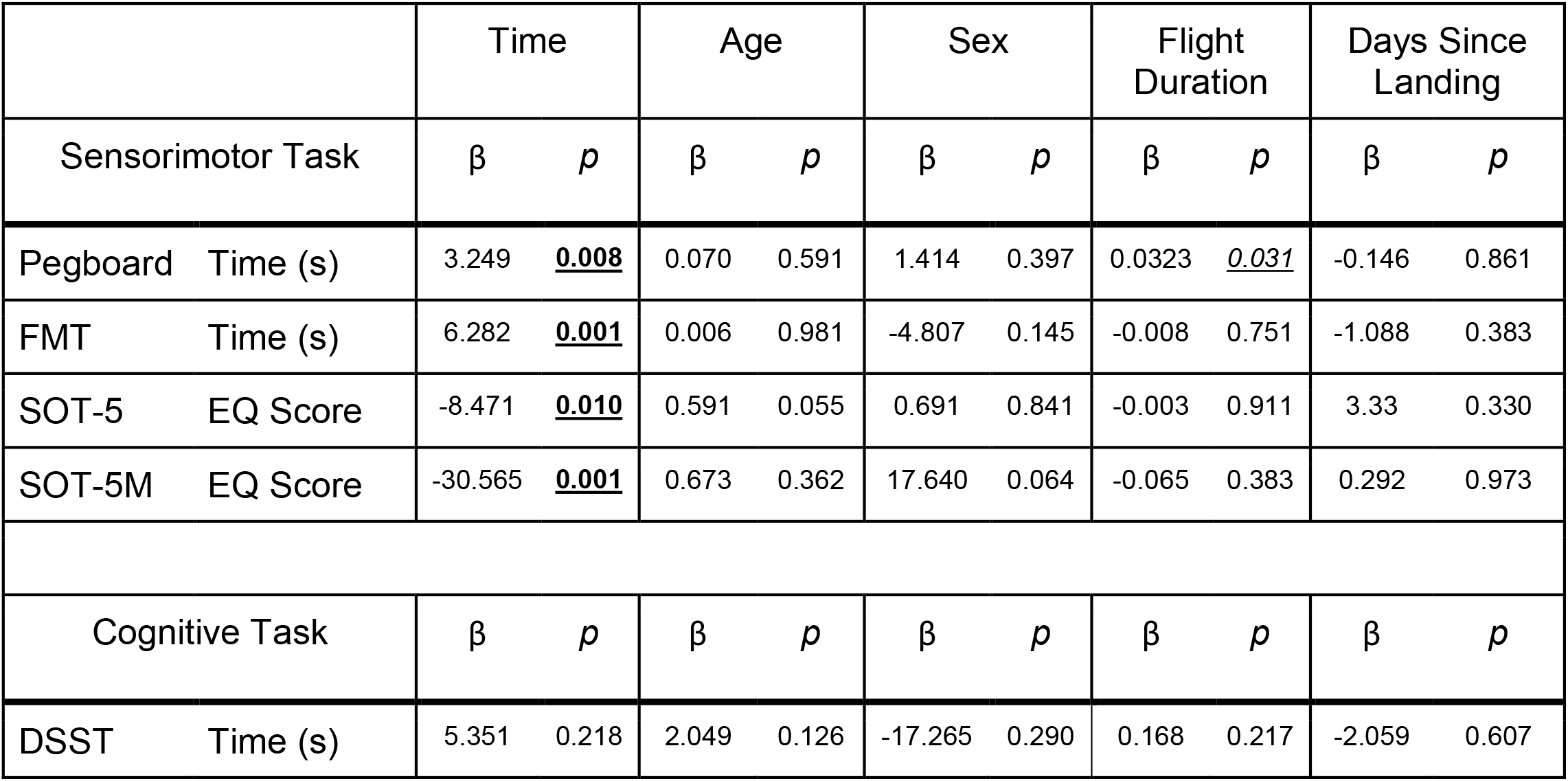
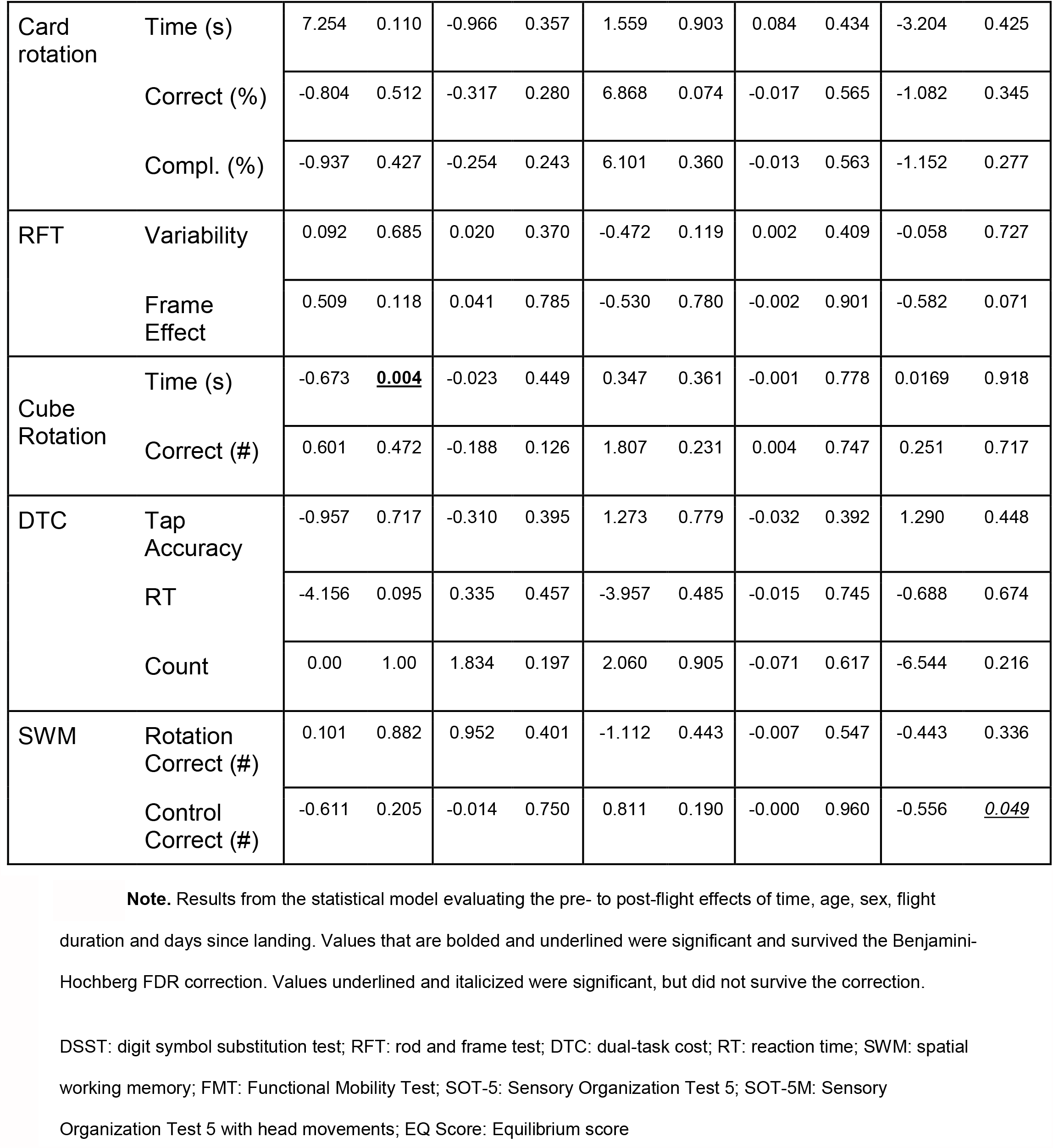
Effects of the microgravity environment

|  | Time |  | Age |  | Sex |  | Flight Duration |  | Days Since Landing |  |
| --- | --- | --- | --- | --- | --- | --- | --- | --- | --- | --- |
| Sensorimotor Task | $\beta$ | $p$ | $\beta$ | $p$ | $\beta$ | $p$ | $\beta$ | $p$ | $\beta$ | $p$ |
| Pegboard Time (s) | 3.249 | **<u>0.008</u>** | 0.070 | 0.591 | 1.414 | 0.397 | 0.0323 | **<u>0.031</u>** | -0.146 | 0.861 |
| FMT Time (s) | 6.282 | **<u>0.001</u>** | 0.006 | 0.981 | -4.807 | 0.145 | -0.008 | 0.751 | -1.088 | 0.383 |
| SOT-5 EQ Score | -8.471 | **<u>0.010</u>** | 0.591 | 0.055 | 0.691 | 0.841 | -0.003 | 0.911 | 3.33 | 0.330 |
| SOT-5M EQ Score | -30.565 | **<u>0.001</u>** | 0.673 | 0.362 | 17.640 | 0.064 | -0.065 | 0.383 | 0.292 | 0.973 |
| Cognitive Task | $\beta$ | $p$ | $\beta$ | $p$ | $\beta$ | $p$ | $\beta$ | $p$ | $\beta$ | $p$ |
| DSST Time (s) | 5.351 | 0.218 | 2.049 | 0.126 | -17.265 | 0.290 | 0.168 | 0.217 | -2.059 | 0.607 |

**Table 2 Note.** Results from the statistical model evaluating the pre- to post-flight effects of time, age, sex, flight duration and days since landing.
|  |  |  |  |  |  |  |  |  |  |  |  |
| --- | --- | --- | --- | --- | --- | --- | --- | --- | --- | --- | --- |
| Card rotation | Time (s) | 7.254 | 0.110 | -0.966 | 0.357 | 1.559 | 0.903 | 0.084 | 0.434 | -3.204 | 0.425 |
|  | Correct (%) | -0.804 | 0.512 | -0.317 | 0.280 | 6.868 | 0.074 | -0.017 | 0.565 | -1.082 | 0.345 |
|  | Compl. (%) | -0.937 | 0.427 | -0.254 | 0.243 | 6.101 | 0.360 | -0.013 | 0.563 | -1.152 | 0.277 |
| RFT | Variability | 0.092 | 0.685 | 0.020 | 0.370 | -0.472 | 0.119 | 0.002 | 0.409 | -0.058 | 0.727 |
|  | Frame Effect | 0.509 | 0.118 | 0.041 | 0.785 | -0.530 | 0.780 | -0.002 | 0.901 | -0.582 | 0.071 |
| Cube Rotation | Time (s) | -0.673 | **<u>0.004</u>** | -0.023 | 0.449 | 0.347 | 0.361 | -0.001 | 0.778 | 0.0169 | 0.918 |
|  | Correct (#) | 0.601 | 0.472 | -0.188 | 0.126 | 1.807 | 0.231 | 0.004 | 0.747 | 0.251 | 0.717 |
| DTC | Tap Accuracy | -0.957 | 0.717 | -0.310 | 0.395 | 1.273 | 0.779 | -0.032 | 0.392 | 1.290 | 0.448 |
|  | RT | -4.156 | 0.095 | 0.335 | 0.457 | -3.957 | 0.485 | -0.015 | 0.745 | -0.688 | 0.674 |
|  | Count | 0.00 | 1.00 | 1.834 | 0.197 | 2.060 | 0.905 | -0.071 | 0.617 | -6.544 | 0.216 |
| SWM | Rotation Correct (#) | 0.101 | 0.882 | 0.952 | 0.401 | -1.112 | 0.443 | -0.007 | 0.547 | -0.443 | 0.336 |
|  | Control Correct (#) | -0.611 | 0.205 | -0.014 | 0.750 | 0.811 | 0.190 | -0.000 | 0.960 | -0.556 | **<u>0.049</u>** |

DSST: digit symbol substitution test; RFT: rod and frame test; DTC: dual-task cost; RT: reaction time; SWM: spatial working memory; FMT: Functional Mobility Test; SOT-5: Sensory Organization Test 5; SOT-5M: Sensory Organization Test 5 with head movements; EQ Score: Equilibrium score

**Table 3:** Recovery from the microgravity environment

Note. Results from the statistical model evaluating the recovery from spaceflight effects of days returned, age, sex and flight duration.
|  | Days Since Return |  | Age |  | Sex |  | Flight Duration |  |
| --- | --- | --- | --- | --- | --- | --- | --- | --- |
| Sensorimotor Task | $\beta$ | $p$ | $\beta$ | $p$ | $\beta$ | $p$ | $\beta$ | $p$ |
| Pegboard Time (s) | -0.136 | **<u>0.016</u>** | 0.176 | 0.274 | 0.748 | 0.716 | 0.020 | 0.248 |
| FMT Time (s) | -0.030 | **<u>0.0001</u>** | 0.031 | 0.877 | -4.196 | 0.121 | 0.007 | 0.753 |
| SOT-5 EQ Score | 0.019 | 0.063 | 0.419 | 0.055 | 1.505 | 0.584 | -0.007 | 0.748 |
| SOT-5M EQ Score | 0.106 | **<u>0.005</u>** | 0.560 | 0.177 | 9.853 | 0.078 | -0.006 | 0.884 |
| Cognitive Task | $\beta$ | $p$ | $\beta$ | $p$ | $\beta$ | $p$ | $\beta$ | $p$ |
| Cube Rotation Time (s) | -0.001 | 0.331 | -0.024 | 0.319 | 0.489 | 0.125 | 0.001 | 0.672 |

FMT: Functional Mobility Test; SOT-5: Sensory Organization Test 5; SOT-5M: Sensory Organization Test 5 with head movements; EQ Score: Equilibrium score

**Table 4:** Direct effects of the microgravity environment

Note. Here we present the results from the statistical models testing for performance changes from pre- to in-flight, controlling for age at launch and sex. In this case, no models yielded statistically significant results.
|  |  | Days Since Launch |  | Age |  | Sex |  |
| --- | --- | --- | --- | --- | --- | --- | --- |
| Task | Measure | $\beta$ | $p$ | $\beta$ | $p$ | $\beta$ | $p$ |
| DTC | Tap | -0.035 | 0.828 | -0.548 | 0.149 | 5.333 | 0.291 |
|  | RT | -0.071 | 0.744 | 0.531 | 0.400 | -2.268 | 0.789 |
|  | Count | 0.320 | 0.580 | 1.620 | 0.136 | 9.600 | 0.500 |

Abbreviations: DTC: dual task cost; RT: reaction time

**Table 5:**
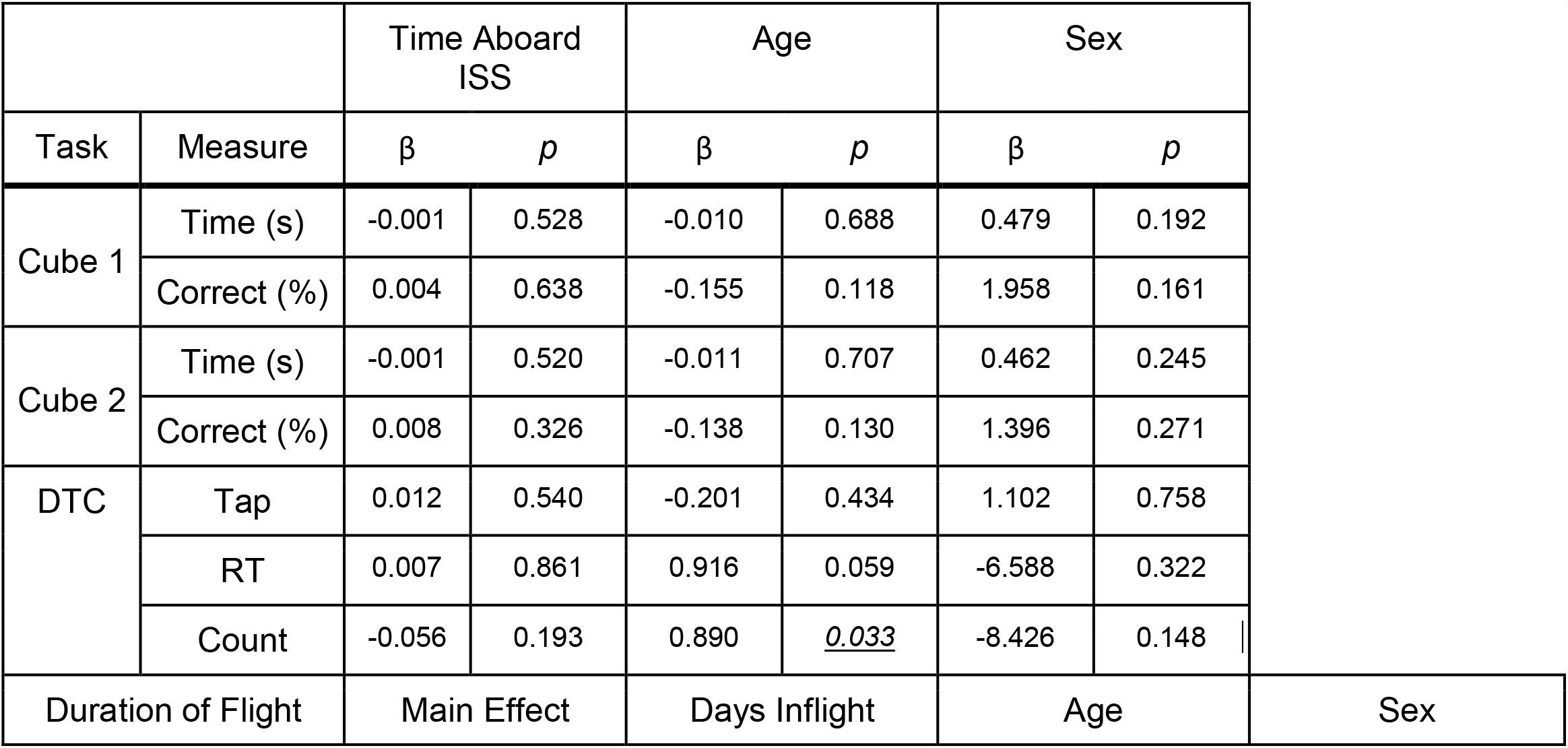
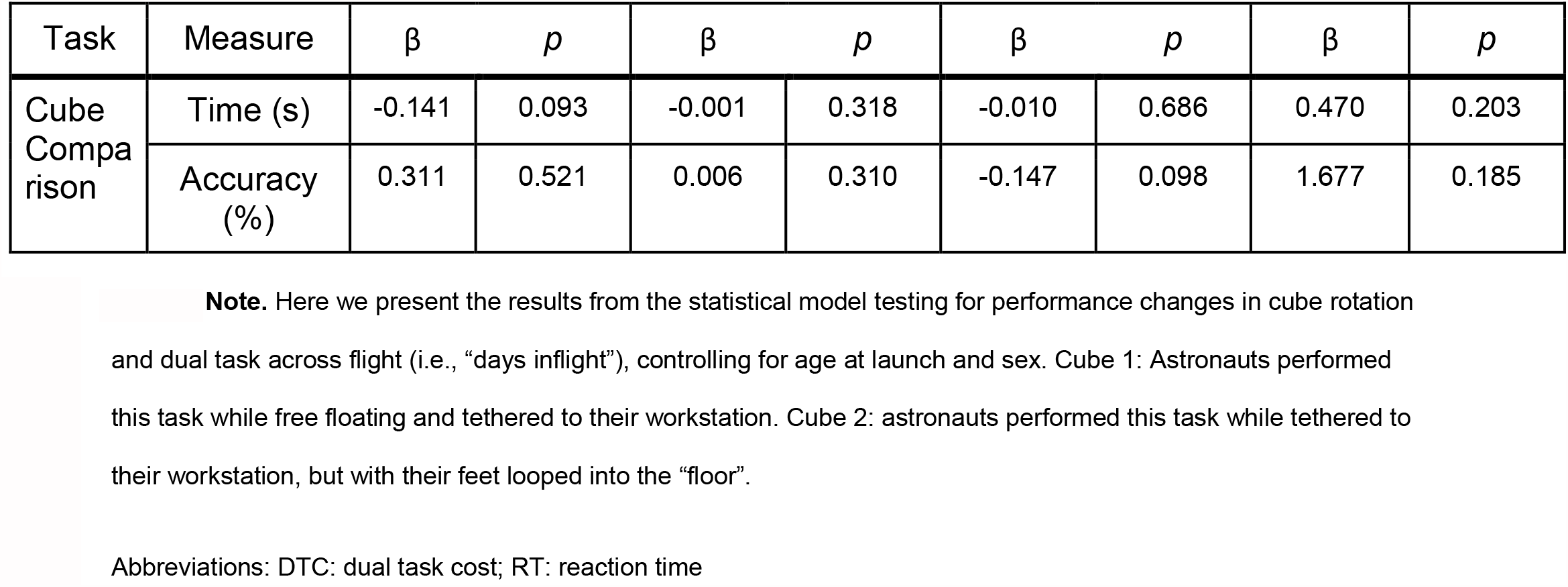
Effects of duration aboard the ISS

##### Model 1) The effect of the microgravity environment

In this model, we considered time as a (fixed effect) categorical variable (pre-flight versus post-flight). We were primarily interested in the statistical significance of this categorical variable (i.e. whether any pre-flight to post-flight changes in performance occurred). We adjusted for the timing variability of the first post-flight session day (R+1 or R+4) by including the (mean-centered) time between landing and the first post-flight session as a covariate, as re-adaptation likely begins as soon as astronauts return to Earth. Mean centered age at launch, sex, and total flight duration were also entered into the model as covariates.

##### Model 2) Recovery from the microgravity environment

This model was only applied for measures where we observed significant changes from pre- to post-flight in model 1, in order to assess post-flight re-adaptation. Here, the fixed effect of time was considered as a continuous variable; we were primarily interested in whether there was a significant effect of time across these post-flight session, to assess the post-flight recovery profile. As in model 1, mean centered age at launch, sex, and total flight duration as covariates.

##### Model 3) Direct effects of microgravity

This model only measured in-flight performance. We utilized time as a continuous variable to evaluate performance changes from pre-flight (L-60) to the first in-flight time point (FD30). Only the in-flight metrics (cube rotation, dual-tasking) were included in this analysis. Mean centered age and sex were included as covariates.

##### Model 4) Effects of duration aboard the ISS

This model only measured in-flight performance for the duration of the mission. We utilized time as a continuous variable to evaluate changes in performance across the three testing periods during spaceflight (FD30, FD90 and FD150). Mean centered age and sex were included as covariates. Since conditions for Cube 2 could only be replicated in spaceflight, we tested for a main effect (cube 1 vs cube 2) for this task.

## 3. Results

Tables 2-5 present all results from the statistical models. Bolded and underlined results remained significant at FDR<0.05. Italicized and underlined results were significant before FDR correction, but did not remain significant following FDR correction.

### 1) The effect of the microgravity environment

We identified significant pre-flight to post-flight performance declines in all sensorimotor tasks (Table 2). FMT completion time increased from pre- to post-flight (*p*=0.001, Fig. 2) as astronauts were slower post-flight. We also identified significant pre-flight to post-flight balance declines, reflected as Equilibrium Quotient scores decreased on both the SOT-5 (*p*=0.011, Fig. 3), and SOT-5M (*p*=0.003, Fig. 4). Astronauts also had a significant increase in completion time on the bimanual Purdue Pegboard Test; that is they were slower to complete the task post-flight (*p*=0.007, Fig. 5).

**Figure 2.**
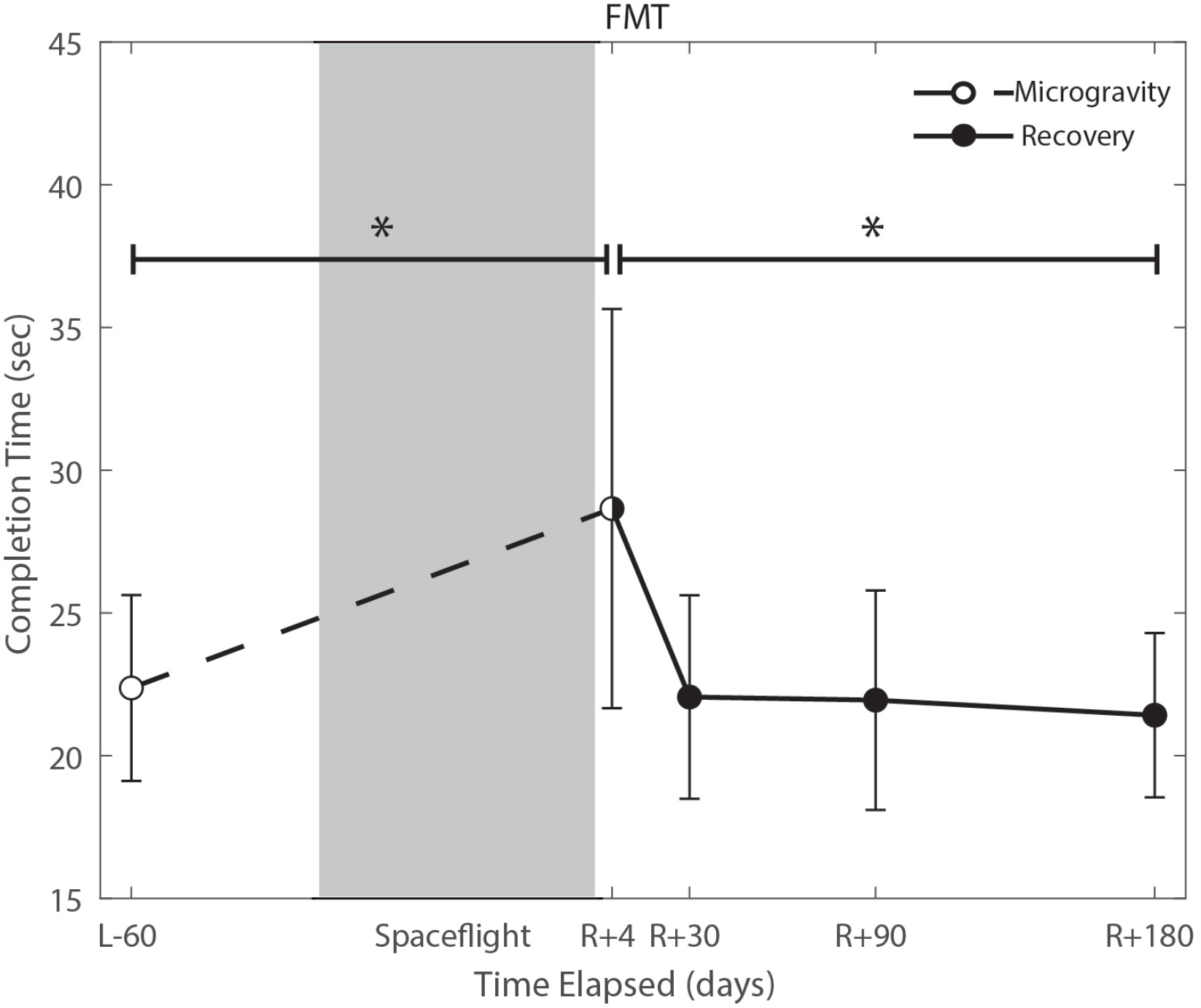
Functional Mobility Test (FMT) performance changes from pre- to post-flight spaceflight and post-flight recovery. Spaceflight resulted in a significant decrease in completion time (seconds; p=0.001). Completion time recovered to baseline levels by approximately 30 days post-flight (p=0.0001).

**Figure 3.**
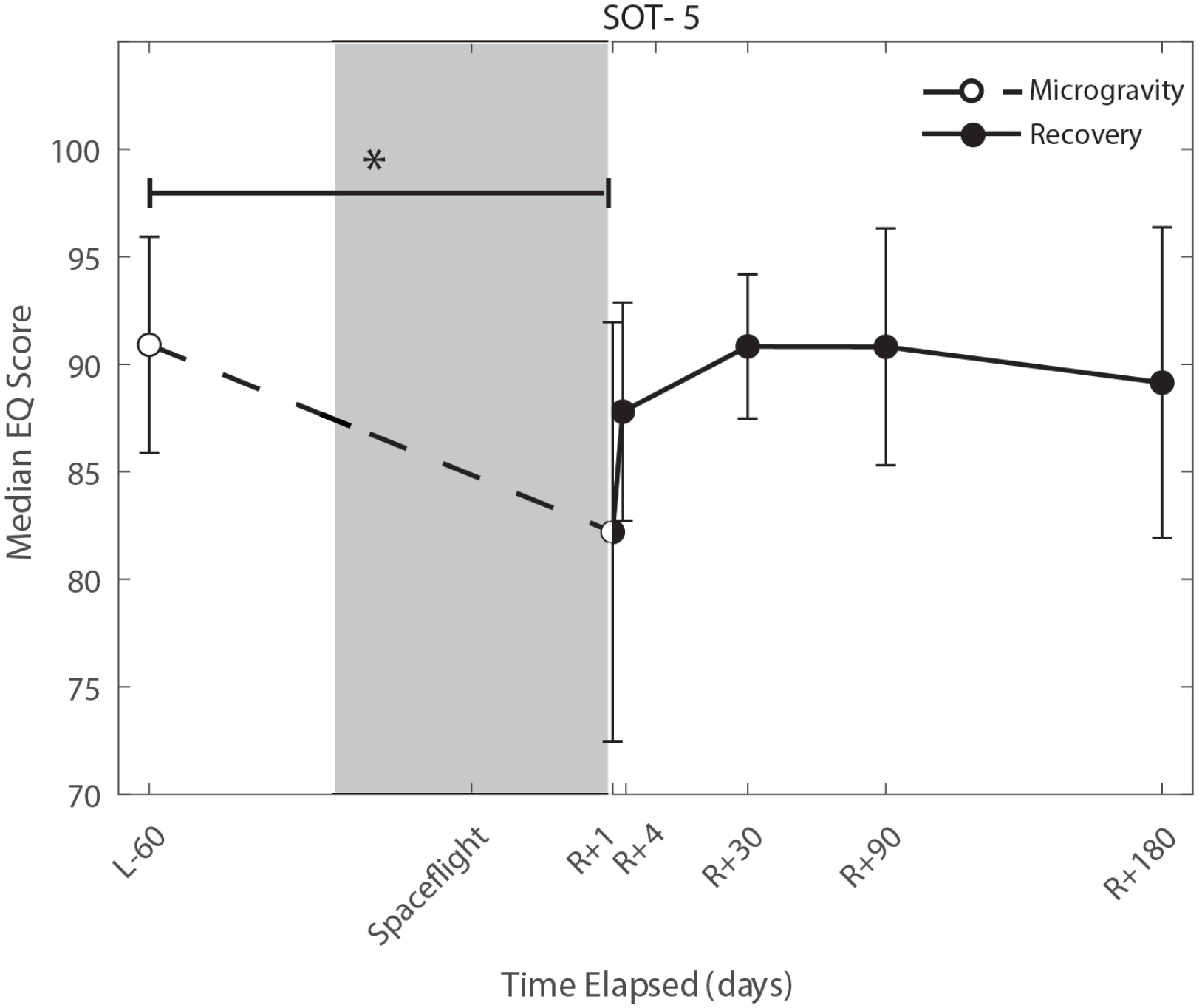
Balance (SOT-5) changes from pre- to post-flight and post-flight recovery. The Sensory Organization Task 5 (SOT-5) performance changes indicate an effect of the microgravity environment resulted in a significant decrease in Equilibrium Score (p=0.01), that did not show statistically significant recovery.

**Figure 4.**
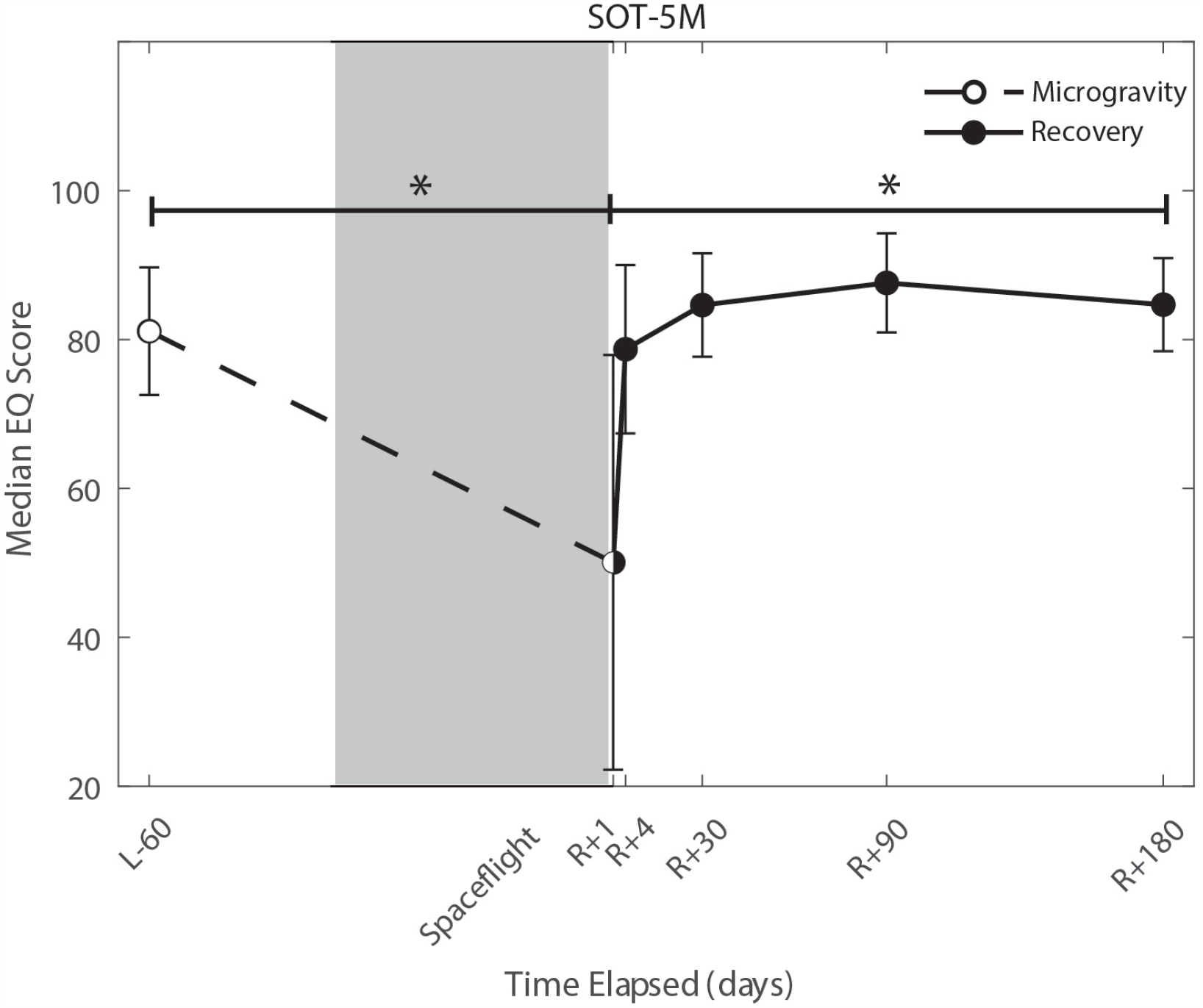
Balance (SOT-5M) changes from pre- to post-flight and post-flight recovery. Sensory Organization Task 5 with head movements (SOT-5M) performance changes indicate an effect of the microgravity environment resulted in a significant decrease in Equilibrium Score (p=0.001). There was a significant recovery of performance following spaceflight (p=0.005).

**Figure 5.**
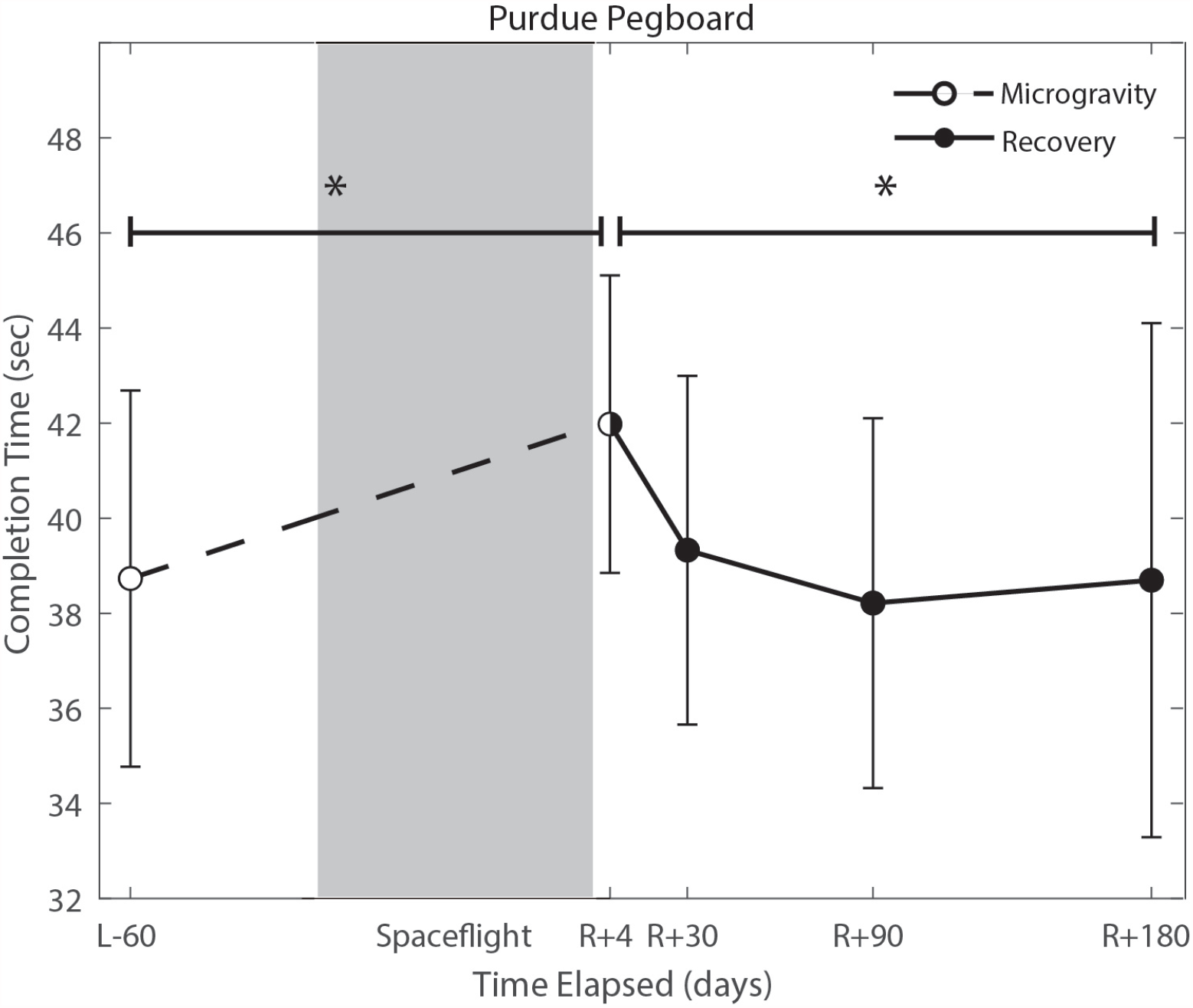
Bimanual Purdue Pegboard completion time changes from pre- to post-flight and post-flight recovery. There is a significant increase in completion time (p=0.008) pre- to post-flight. There is a significant change in recovery (p=0.016).

With the exception of cube rotation, no cognitive assessments showed pre-flight to post-flight changes. Cube rotation response time decreased significantly post-flight (*p*=0.004; Fig. 6); as astronauts showed faster cube rotation completion time post-flight. We also idenitifed a significant effect of days since landing on the SWM control task (*p*=0.049); where a longer time delay between landing and the first session was associated with better SWM control task performance.

**Figure 6.**
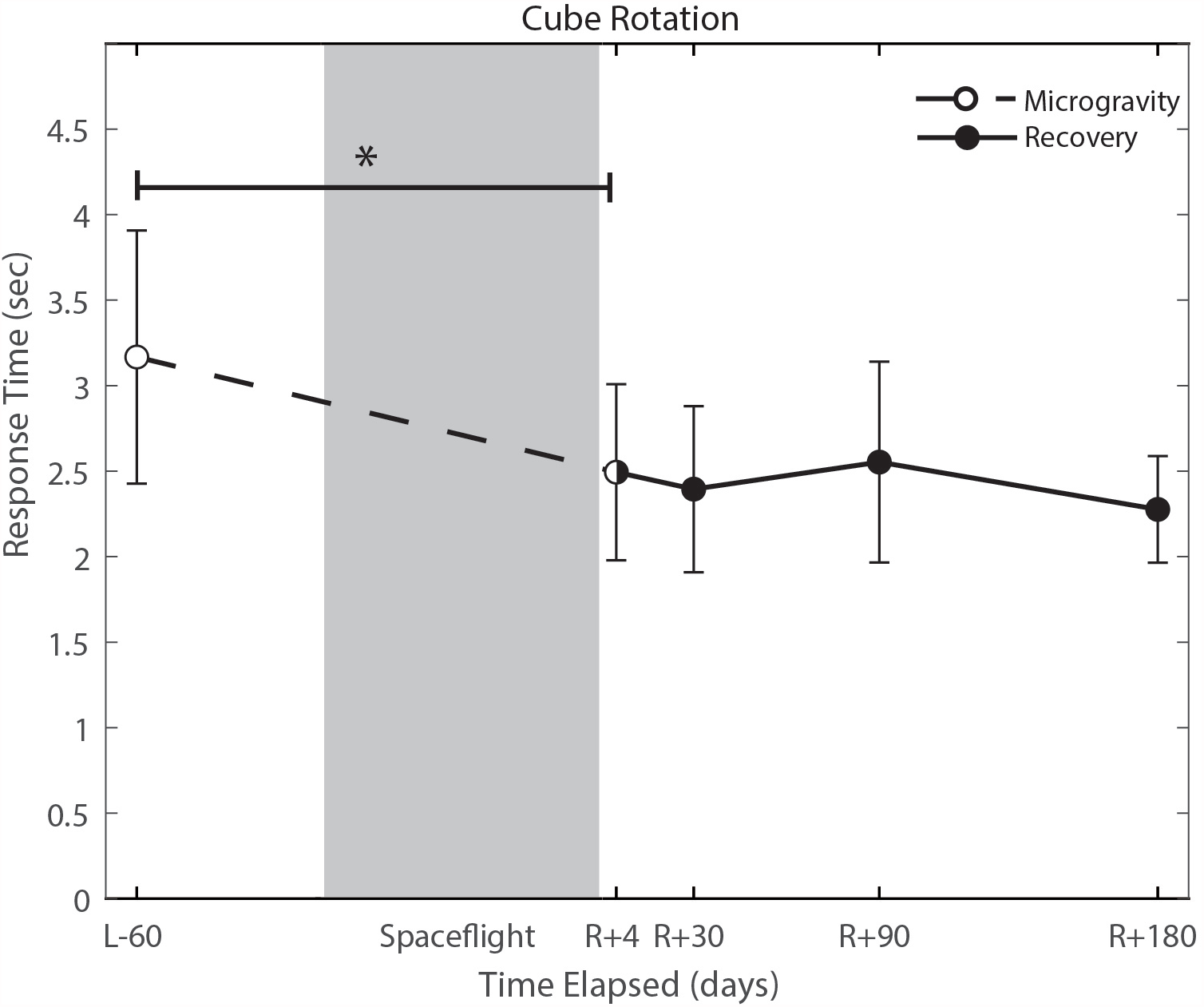
Cube rotation performance changes from pre- to post-flight and post-flight recovery. Subject’s response time improved (p=0.004) as a significant decrease in response time.

### 2) Recovery from the microgravity environment

Of the measures that changed significantly from pre-flight to post-flight, we observed significant post-flight recovery (Table 3) on the bimanual Purdue Pegboard Test (*p*=0.0158), FMT (*p*=0.0001) and the SOT-5M (*p*=0.0062). Astronauts’ performance on the bimanual Purdue Pegboard Test returned to near baseline levels by 30 days post-flight and continued to improve by 90 days post-flight (Fig. 5). FMT performance showed similar trends, with a return to pre-flight performance levels by R+30 (Fig. 2). SOT-5M scores showed substantial improvements in performance from R+1 to R+4 that continued to improve at R+30 and R+90 before plateauing (Fig. 4).

### 4) Direct effects of microgravity

Astronauts performed two cognitive tasks (cube rotation and dual-tasking) aboard the ISS, first approximately 30 days after their arrival. There was no significant pre- to in-flight performance changes on these tasks (Table 4).

### 5) Effects of duration on the ISS

There were no significant changes in performance of the cube rotation or dual-tasking assessments across the three in-flight time points (Table 5).

## 4. Discussion

The current study was designed to investigate sensorimotor and cognitive performance changes associated with long-duration spaceflight and their subsequent recovery post-flight.

Consistent with previous results (Reschke et al., 1994a; 1994b; 1998; Mcdonald et al. 1996, Bloomberg et al. 1997; Layne et al. 1998; Mulavara et al, 2010; Mulavara et al., 2018; Miller et al., 2018), we found pre- to post-flight declines in balance and mobility. There were also declines in bimanual coordination from pre- to post-flight, as indicated by performance on the bimanual Purdue Pegboard Test. All of these measures were shown to recover by 30 days after return to Earth. There were no significant effects of spaceflight on the cognitive measures collected here, including pre- to post-flight and pre- to in-flight performance comparisons.

### 4.1. Whole-body postural and locomotor control

Sensorimotor deficits due to spaceflight have been previously reported following both short (weeks) and long (months) duration spaceflight (Reschke et al., 1994a; 1994b; 1995; 1998; Mcdonald et al. 1996, Bloomberg et al. 1997; Layne et al. 1998; Mulavara et al, 2010; Mulavara et al., 2018; Miller et al., 2018); here, we find similar declines and subsequent recovery profiles in locomotion and balance. These balance and gait findings support the argument that adaptive sensory reweighting occurs during spaceflight. While in the microgravity environment of space, the vestibular system is limited to receive acceleration inputs from the otoliths. The central nervous system adapts by upweighting other sensory inputs (e.g. visual and proprioceptive inputs). Upon return to Earth, vestibular afferent inputs return to normal levels and in-flight adaptations become maladaptive. As evident in Figures 3 and 4, SOT-5 and SOT-5M performance show significant deficits at R+1; however, by R+4 postural control has returned to near baseline levels. There appear to be some slow, persisting effects out to R+30, suggesting both rapid and slower re-adaptation processes. Adaptation of reaching movements to visuomotor conflict (e.g., visuomotor rotation where visual feedback is offset as a perturbation) on Earth has been well-studied. This literature suggests that early adaptive changes are more cognitive and strategic in nature whereas slower changes reflect more implicit, procedural processes (Anguera et al., 2010; Taylor & Ivry, 2013; McDougle et al., 2015; Christou et al. 2016). It is unclear whether similar processes are at work when adapting to sensory conflict on Earth and adapting to the sensory conflict created by microgravity, but the initial fast recovery followed by a slower timeline to reach pre-flight levels suggests the possibility of similar processes.

### 4.2. Fine motor control

Novel findings here include a significant increase in bimanual Purdue Pegboard Test completion time. We fit a linear regression model between age and bimanual Purdue Pegboard completion time in a control sample of 24 subjects (mean age 33.292, 8 female), and found that completion time increased by 0.13 seconds per year of age. The reported increase of 3.25 seconds exhibited by crewmembers is approximately equivalent to a 25 year age difference. However, it should be noted that the controls were, on average, 14 years younger than astronaut crewmembers, which may result in overestimation of years decline pre to postflight. Previous reports of fine motor control declines following spaceflight include impairments in force modulation (Rafiq et al., 2006), surgical operating completion time (Campbell et al., 2005), keyed pegboard completion time (Mulavara et al., 2018), decreased unimanual Purdue Pegboard performance (Moore et al., 2019), reaction time, movement duration, and response amplitude (Bock et al., 2003). These findings have raised concerns that astronauts will face increased risk of operational task failure (Paloski et al., 2008). The results from the current study further support previous findings that there is a marked impairment in fine motor control due to spaceflight, including bimanual coordination. Moreover, these changes are evident up until 30 days post-flight. While the mechanisms underlying these manual motor control declines are unclear, it has been shown that there is an increase in skin sensitivity for both fast and slow receptors following spaceflight (Lowrey et al., 2014). This upweighting of tactile inputs may be adaptive inflight when the body is unloaded, but could potentially be maladaptive upon return to Earth, resulting in these transient manual motor performance declines.

### 4.3. Cognitive measures

Cognitive declines with spaceflight have not conclusively been observed. Changes that have been reported include an increased ability to mentally rotate their environment, and decreased ability to spatially orient letters in a word during early short duration spaceflight (Clément et al., 1987), reduced cognitive-motor dual-tasking ability (Manzey et al., 1995; 1998; Bock et al., 2010), increases in risky behavior in a single subject case study (Garrett-Bakelman et al., 2019), and anecdotal reports of “space fog” (Clément et al., 2020). In the present study, we investigated a range of cognitive domains both from pre- to post-flight and while astronauts were aboard the ISS. The only significant changes from pre- to post-flight that survived FDR correction was in the cube rotation response time, which showed a decrease in response time that is likely attributable to a practice effect. Astronauts performed the cube rotation task twice per test session aboard the ISS, once while free floating yet tethered to the laptop console and again while tethered with feet in loops on the “floor”. These two setups allowed us to identify whether somatosensory feedback associated with having the feet on the “floor” and performing the task in a “seated” posture provides spatial orientation cues to aid in mental rotation performance. There were no statistically significant differences (*p*=0.093) between cube 1 (feet unattached) and cube 2 (feet attached), however there were trend level effects of a faster response time on cube 2. These results may be limited by our small sample size, but could potentially have operational relevance. This trend level effect could reflect practice.

### 4.4. Mission duration

A current focus in spaceflight research is understanding the effects of flight duration on the human brain and behavior. NASA is planning to return to the Moon with the Artemis program and Mars by the 2030’s. A round trip to Mars is estimated to be around 21 months, which is longer than any current astronaut has spent in space on any given mission. This makes it imperative to understand whether there is a “dose-dependent” effect of spaceflight stressors/hazards on human performance. In the current study, most astronauts had mission durations of approximately 6 months, but there was a range with some crewmembers spending nearly 12 months (ranging from 4 to 11 months in space). We included mission duration in our statistical models to investigate its effect, finding only an uncorrected decline in bimanual Purdue Pegboard Test completion time with longer flight duration. The lack of spaceflight duration effects on our results may suggest that there are little functional changes associated with mission duration; however, it has been shown recently that the magnitude of spaceflight-associated structural brain changes is directly related to mission duration. We (Hupfeld et al. 2020a) recently reported that astronauts who spent one year in space exhibited larger magnitude brain fluid shifts, greater right precentral gyrus gray matter volume and cortical thickness changes, greater supplementary motor area gray matter volume changes, and greater free water volume changes within the frontal pole. Six-month missions were shown to result in greater increases in cerebellar volume as compared to 12-month missions. Brain changes exhibited only partial recovery at six months post-flight (Hupfeld et al. 2020a). Our work and other studies have also reported that there are persisting ventricular volume changes at six months and one year post-flight (Hupfeld et al. 2020a; Van Ombergen et al., 2019; Kramer et al., 2020; Jillings et al., 2020). It is important to consider these brain changes; it is possible that behavior has returned to pre-flight levels by one month post-flight without a concomitant return to pre-flight neural control patterns. That is, there may be a substitution of brain networks or compensation that is still taking place post-flight even when behavior has recovered, without restitution of pre-flight brain (Rothi et al. 1983, Hupfeld et al., 2020b).

This study is one of the few to have collected longitudinal data from astronauts on the ISS, allowing us to directly examine the effects of initial and longer term microgravity exposure. One of the tasks measured during spaceflight required single and dual-tasking. Dual-tasking has been evaluated previously during spaceflight; results showed impairments in both cognitive and motor behaviors in long duration spaceflight missions (Manzey et al. 1995; 1998), with dual-task costs greater in space than on Earth. Additionally, these impairments were greatest during early flight and stabilized after approximately 9 months in space. However, these two reports were single subject case studies. Bock et al. (2010) further investigated dual-tasking in microgravity with a larger cohort of 3 astronauts performing a tracking task while also performing one of four reaction time tasks. They found an overall increase in tracking error and reaction time under dual-task conditions. The present results differ as we found no differences in dual-task costs upon arrival to the ISS (performance measured at approximately 30 days into the flight and compared to pre-flight), nor as flight duration increased (performance measured at approximately 90 and 180 days into the flight). This may be due to a difference in complexity of the cognitive and motor tasks, a difference in the underlying task mechanisms, or due to the larger sample evaluated here.

### 4.5. Limitations

One of the primary limitations of this study is the small number of female astronauts; of the fifteen participants, only three were female. This does not allow sufficient power to evaluate sex differences. Another limitation in this study is the time delay between landing and the initial post-flight data collection, as astronauts re-adapt to Earth’s gravity relatively quickly. We found that postural control returned to baseline levels within roughly four days post-flight. It is possible that some of our other measures respond in a similar manner; this would mean that, by post-flight day four, we may have missed many spaceflight-related performance changes. Moreover, we did not have test sessions between post-flight days 4 and 30, limiting our ability to delineate post-flight rapid recovery curves.

In this study, we evaluated the effects of the microgravity environment on astronauts’ sensorimotor and cognitive performance with a range of behavioral measures collected before, during, and following missions to the ISS. We found marked decreases in balance, mobility and bimanual coordination following exposure to the microgravity environment. These declines are transient and return to baseline levels within roughly 30 days. Additionally, we identified a trend for increased cognitive performance on some measures when astronauts had their feet on the “floor” of the ISS, suggesting that additional orientation cues may increase spatial working memory ability in microgravity. In the same sample, we also collected functional MRI data during task performance before and following spaceflight as well as measures of brain structure (structural MRI and diffusion weighted MRI). In future analyses, we will examine brain changes and their relation to behavioral performance. It may be that, in cases where we do not see behavioral changes, the underlying networks engaged for task performance will have changed in a compensatory fashion due to spaceflight. Further analyses of our neuroimaging data in conjunction with these performance measures will give us insight into the adaptive or maladaptive effects of spaceflight.

## Acknowledgements

This work was supported by grants from the National Aeronautics and Space Administration (NASA NNX11AR02G to R.S., A.M., S.W., P.A.R.L. and J.B. During the completion of this work G.T. was supported by the University of Florida’s (UF) Graduate Student Funding Award. K.H. was supported by a National Science Foundation Graduate Research Fellowship under Grant no. DGE-1315138 and DGE-1842473, National Institute of Neurological Disorders and Stroke training grant T32-NS082128, and National Institute on Aging fellowship 1F99AG068440. H.M. was supported by a Natural Sciences and Engineering Research Council of Canada (NSERC) Postdoctoral Fellowship and a NASA Human Research Program Augmentation Grant. The authors would like to thank all of the astronauts who volunteered their time, without them this project would not have been possible.

